# Deconvolution of the on-target activity of plasmepsin V peptidomimetics in *P. falciparum* parasites

**DOI:** 10.1101/2025.07.24.666510

**Authors:** Wenyin Su, William Nguyen, Ghizal Siddiqui, Jerzy M. Dziekan, Danushka Marapana, Jocelyn S. Penington, Somya Mehra, Zahra Razook, Kirsty McCann, Anna Ngo, Kate E. Jarman, Alyssa Barry, Anthony T. Papenfuss, Paul R. Gilson, Darren J. Creek, Alan F. Cowman, Brad E. Sleebs, Madeline G. Dans

## Abstract

Plasmepsin V (PMV), an essential aspartyl protease, plays a critical role during the asexual blood stage of infection of *Plasmodium* by enabling the export of parasite proteins into the host red blood cell. This export is vital for parasite survival and pathogenesis, making PMV an attractive target for antimalarial drug development. Peptidomimetic inhibitors designed to mimic the natural substrate of PMV have demonstrated potent parasite-killing activity by blocking protein export. While these compounds have been instrumental in validating PMV as a bona fide antimalarial target, inconsistencies between their biochemical potency and cellular activity have raised questions regarding their precise mechanism of action. In this study, we employed chemoproteomic approaches, including solvent induced protein precipitation (SIP) and intact-cell thermal PISA profiling, to demonstrate PMV target engagement by the peptidomimetics. To further support these findings, we generated parasite lines exhibiting reduced sensitivity to peptidomimetics. Through whole-genome sequencing of these parasite lines, a single nucleotide variant (SNV) within the *pmv* gene was revealed. This mutation was later validated using reverse genetics, confirming its role in mediating resistance. Together, these data provide strong evidence that the peptidomimetics exert their antimalarial activity by directly targeting PMV. These findings further support the potential of PMV as a validated and promising target for future antimalarial drug development.

## INTRODUCTION

There is an urgent need to develop new strategies and agents to treat malaria because of the emergence of resistance against antimalarial therapies in clinical use [1, 2]. This is particularly significant given that malaria continues to pose a substantial global health burden, with an estimated 263 million cases and 597,000 deaths reported in 2023 [2].

*P. falciparum* expresses ten cathepsin-D like aspartyl proteases termed plasmepsins (PM) [3].

PMI-IV are located in the digestive vacuole and are involved in the initial phase of haemoglobin digestion with some functional redundancies [4–6]. PMVI-VIII, whilst required for the transmission stage of the lifecycle, have no essential role in the asexual stage of the lifecycle [3, 7–9]. PMV, PMIX and PMX are all essential in the asexual blood stage of infection; PMIX and PMX are required for parasite egress and invasion of red blood cells (RBCs), and also play a critical role in other stages of the lifecycle [10–12]. Highlighting the druggability of these plasmepsins, inhibitors against PMIX/X are currently in clinical trials (NCT06294912). PMV is essential for protein export and parasite development in both the asexual and sexual stages of the lifecycle [13], and therefore also remains an attractive multi-stage antimalarial target.

During the asexual blood stage, parasites export a large number of proteins to the host RBC which is essential for nutrient and waste exchange, sustenance and to evade the host immune system [14]. Most of these exported proteins contain a pentameric N-terminal motif called the *Plasmodium* export element (PEXEL) [15]. The PEXEL is essential for transportation of proteins from the parasite endoplasmic reticulum (ER), through the parasitophorous vacuole membrane via the *Plasmodium* translocon of exported proteins (PTEX) into the host RBC [16, 17]. The PEXEL is comprised of a highly conserved motif among all *Plasmodium* species and is designated by the consensus sequence, RxLxE/Q/D. The PEXEL motif is recognised and cleaved on the C-terminal side of Leu by the ER resident aspartyl protease, PMV [18]. The remaining C-terminal xE/Q/D stub is then N-acetylated by an acetylase committing the protein for export to the host RBC [19].

One approach to develop a small molecule inhibitor of PMV is to mimic the native PEXEL motif. Historically, aspartyl proteases have been targeted with transition state mimetics that mimic the catalytic intermediate in the proteolysis of the substrate. Examples include peptidomimetics targeting HIV protease, renin and BACE1 [20–22]. The earliest example of a PEXEL transition state peptidomimetic was WEHI-916 (Figure 1). WEHI-916 is a potent inhibitor of PMV and modestly inhibits export of PEXEL containing proteins and parasite development [23, 24]. By replacing the P3 Arg in WEHI-916 with canavanine (Cav), an analogue named WEHI-842 was subsequently produced and has 10-fold improvement in both affinity for PMV and efficacy against *P. falciparum* compared to WEHI-916 (Figure 1) [25, 26]. Further optimisation of the peptidomimetic compounds at the P_2_ position generated two analogues named WEHI-600 and WEHI-601 (Figure 1). Both WEHI-600 and WEHI-601 show 4-fold improvement in both PMV inhibition and parasite efficacy [27].

**Figure 1.**
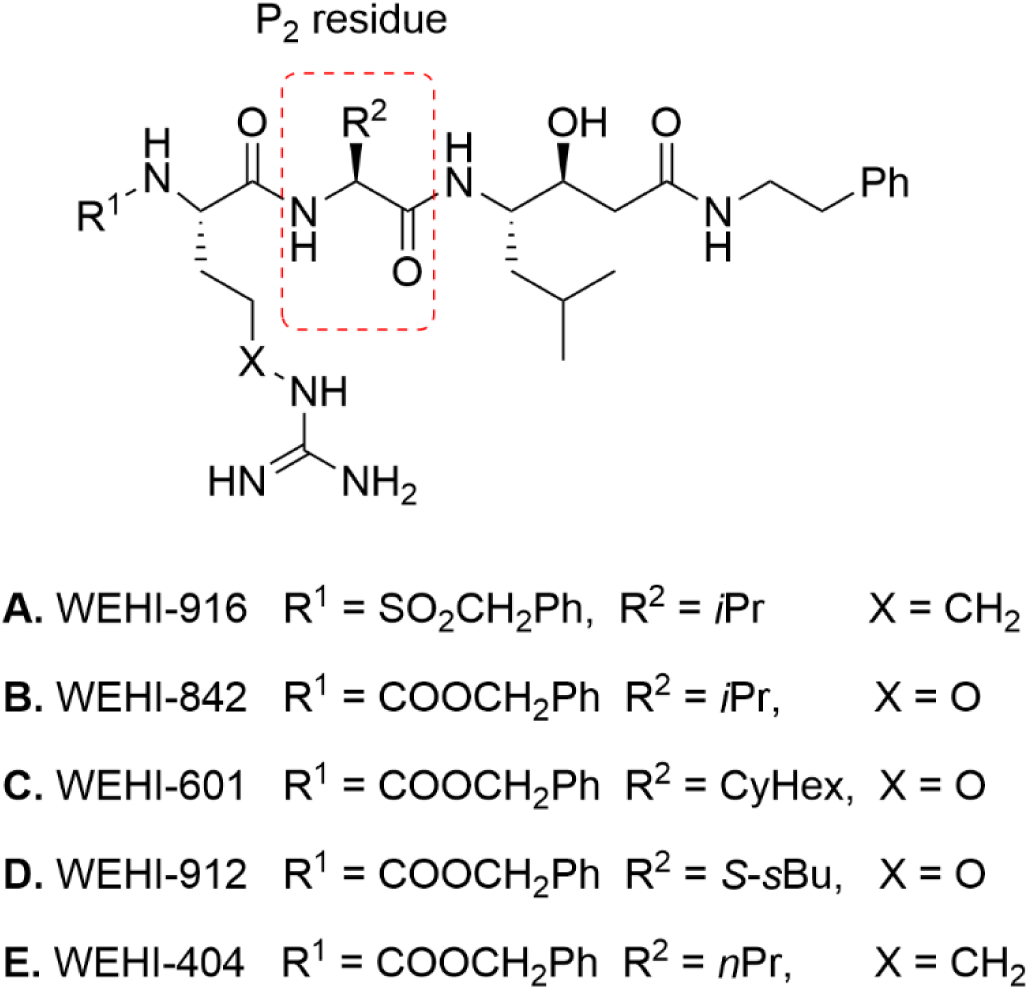
Structure of PMV peptidomimetic inhibitors **(A)** WEHI-916; **(B)** WEHI-842; **(C)** WEHI-601; **(D)** WEHI-912; **(E)** WEHI-404. The P_2_ residue is highlighted in red.

The peptidomimetic inhibitors have been robust chemical probes in the pharmacological validation of PMV. A confounding factor in the development of these inhibitors, is the disparity between the biochemical inhibition of PMV and the concentrations of these peptidomimetics required to block PEXEL cleavage and protein export and kill the parasite. Here we investigated these PMV peptidomimetic inhibitors through chemical and genetic techniques and confirmed their on-target activity against *Plasmodium falciparum*. The observed disparity is likely due to cell permeability and the abundance of PMV expression in the parasite, requiring heightened concentrations of the peptidomimetic to completely nullify its proteolytic function.

## RESULTS

### Plasmepsin and human biochemical selectivity

Given the structural architecture of parasite plasmepsins is similar, it is possible that the peptidomimetics designed to target PMV were also binding other essential plasmepsin proteins, PMIX and PMX, and in turn inhibiting parasite development through one or multiple targets. At the time of the PMV peptidomimetic development, recombinant PMIX and PMX were not accessible [24, 27]. More recently, reliable sources of recombinant PMIX and PMX have become available [12, 28] allowing us to assess the biochemical activity of the PMV peptidomimetics WEHI-601, WEHI-916 and WEHI-842 against these proteins. These assays indicated that the peptidomimetics had no activity (IC_50_ >50 µM) against PMIX, whilst some modest activity against PMX was observed with WEHI-916 and WEHI-842 (IC_50_ 0.99 and IC_50_ 1.85 µM, respectively) but not WEHI-601 (IC_50_ > 99 µM) (Table 1). Despite this activity against PMX, it is notable that WEHI-916 and WEHI-842 were found to inhibit PMV 11- to 62-fold more potently than PMX (Table 1).

**Table 1.** Biochemical activity of peptidomimetics against asexual stage essential plasmepsins and human aspartyl proteases.^a^

| Compound | Parasite<br>EC<sub>50</sub><br>(SD) | PMV<br>IC<sub>50</sub><br>(SD)<sup>^</sup> | PMIX<br>IC<sub>50</sub><br>(SD) | PMX<br>IC<sub>50</sub> (SD) | cathepsin<br>D IC<sub>50</sub><br>(SD) | BACE1<br>IC<sub>50</sub><br>(SD) | renin<br>IC<sub>50</sub><br>(SD) |
| --- | --- | --- | --- | --- | --- | --- | --- |
| **WEHI-916** | 6.05<br>(1.79) | 0.09<br>(0.01) | 50.75<br>(25.58) | 0.99<br>(0.02) | ND | ND | ND |
| **WEHI-842** | 1.07<br>(0.17) | 0.03<br>(0.03) | >99.00 | 1.85<br>(0.24) | 0.52*<br>(0.61) | >1.00 | >1.00 |
| **WEHI-601** | 0.70<br>(0.17) | 0.005<br>(0.002) | >99.00 | >99.00 | ND | >11.10* | ND |

<sup>a</sup>All values are in $\mu\text{M}$ with values derived from at least three biological replicates except for those indicated with a \* which represent two biological replicates. ND indicates not determined.

To assess the peptidomimetic inhibitor’s selectivity, we then evaluated the panel of compounds against human aspartyl proteases. This indicated that the peptidomimetic WEHI-842 does not inhibit the human aspartyl proteases BACE1 and renin (IC_50_ >1.00 µM) (Table 1). WEHI-601 also did not display any activity against BACE1 (IC_50_ >11.0 µM). WEHI-842, however, demonstrated modest activity against human cathepsin D (IC_50_ 0.52 µM). Despite this activity, WEHI-842 has a 16-fold greater activity against PMV (IC_50_ 0.03 µM), indicating selectivity towards the parasite aspartyl protease PMV was maintained.

### Target engagement studies reveal Plasmepsin V as the primary target of WEHI-601

Next, we wanted to investigate the on-target activity of the peptidomimetics against PMV within the parasite. WEHI-601 was selected to carry out these experiments as it displayed the highest potency against both recombinant PMV and *P. falciparum* growth viability. Firstly, we used the principle of Solvent Induced Protein Precipitation (SIP) which reveals protein-ligand interactions based on the alteration of protein susceptibility to denaturation after ligand binding in the presence of an organic solvent [29–31]. We performed SIP assays whereby schizont lysate was treated with 5 µM of WEHI-601, or vehicle solvent, DMSO. We then challenged the lysate with a mixture of 0-25% (v/v) organic solvent acetone/ethanol/formic acid (AEF) and the soluble proteins were isolated by centrifugation. The soluble fractions were separated by western blot and probed with anti-PMV antibody (Figure 2A). Here from a concentration of >17% AEF, we observed significantly more soluble PMV in the WEHI-601 treated group when compared to the DMSO control, indicating target engagement between the peptidomimetic and PMV.

**Figure 2.**
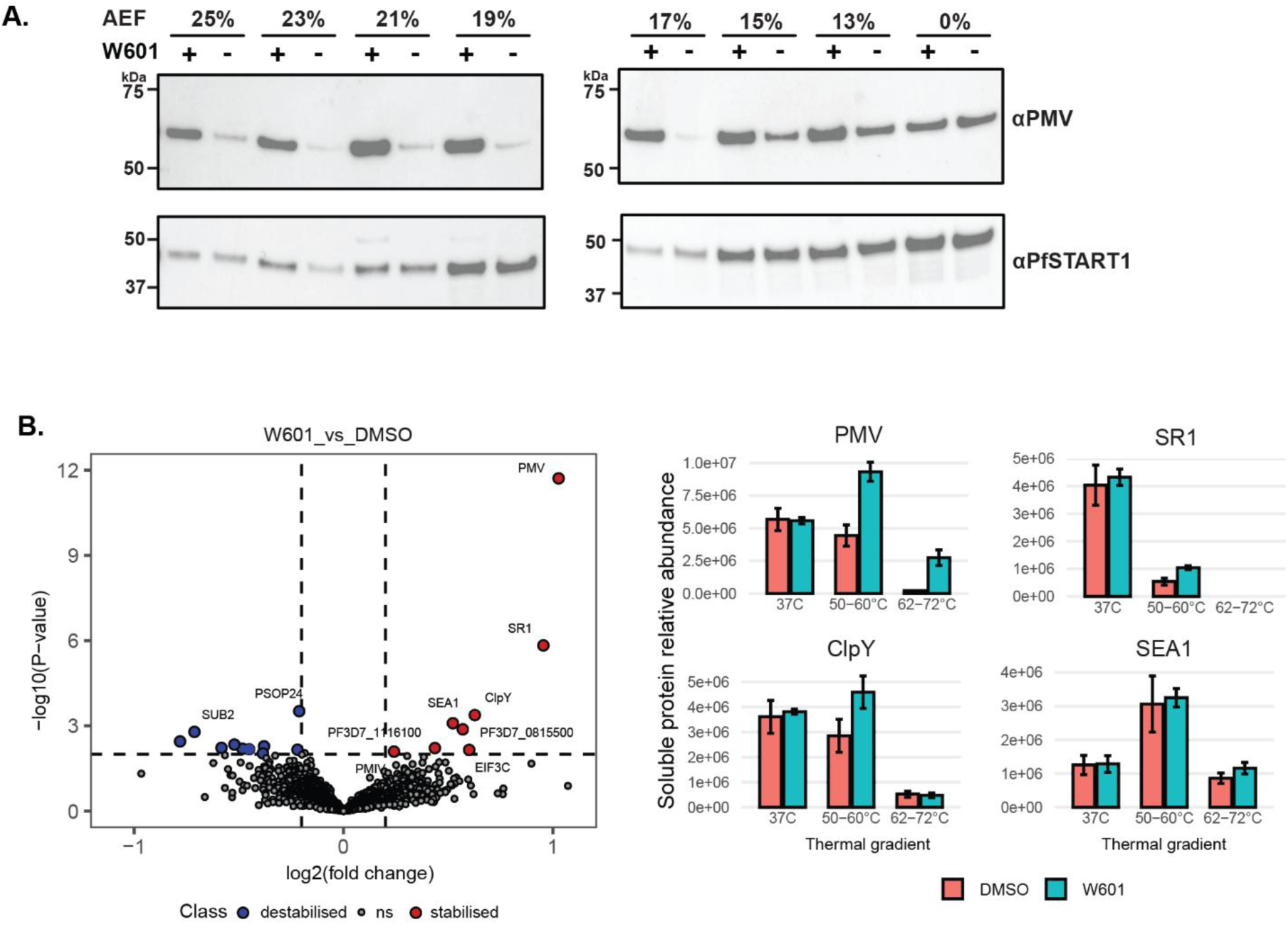
WEHI-601 demonstrates target engagement for plasmepsin V. **(A)** Solvent induced protein precipitation (SIP) assays. The parasite lysate was treated with DMSO (‘-’) or 5 µM WEHI- 601 (‘+’). The lysate was challenged in acetone/ethanol/formic acid mixture (AEF, v:v:v = 50:50:1) gradient 0-25%. The soluble fraction of protein was extracted and soluble proteins were separated out via western blot and probed with anti-PMV antibody. PfSTART1 was probed with anti-PfSTART1 antibody as a loading control. Replicate blots can be found in Figure S1. **(B)** Live cell thermal PISA profiling of WEHI-601 target engagement. Volcano plots depicted differential soluble protein abundance analysis (moderated t-test) for WEHI-601 (10 μM) and DMSO treated parasites following heat pulse heat challenge (n = 4 biological replicates, mean +/− SD). Non-significant (ns) proteins were plotted in grey. Destabilised are plotted in blue and stabilised proteins are plotted in red. Hit selection cuts-off at 0.73 log2 fold change and *p*<0.01 are indicated with dashed lines. The top four significantly stabilised hits are shown in the bar graphs representing the relative soluble protein abundance in WEHI-601 and DMSO treated samples, across three thermal challenge conditions tested (x-axis). *P*-values can be found in Table S1. Bar graphs represent the average value of 4 biological replicates (+/-SD).

We next attempted to investigate any possible engagement between WEHI-601 and PMX in these SIP assays using a haemagglutinin (HA)-tagged PMX parasite line [12]. Despite multiple attempts, we were unable to observe any reproducible pattern of PMX detection via solvent denaturation (Figure S1). This was possibly due to PMX not being compatible with the organic solvent method via western blotting. Thus, to further investigate the selectivity of the peptidomimetic inhibitors in parasites, we conducted a live cell thermal profiling assay coupled with a global proteomics analysis. Following a Proteome Integral Solubility Alteration (PISA) experimental format [32], we separated the samples after thermal challenge and pooled samples into low and high denaturing groups (Gradient 50°-60°C and Gradient 62°-72°C). Through Data Independent Acquisition mass spectrometry (DIA-MS) analysis, relative protein abundance was determined. This assay revealed 18 proteins that were either significantly stabilised or destabilised in the presence of WEHI-601, compared to the DMSO control (Table S1). Amongst these hits, PMV was found to be the most significantly stabilised protein (Figure 2B, p<0.00001). Other proteins that were significantly stabilised included the putative serpentine receptor SR1 (PF3D7_1131100), an ATPase subunit ClpY (PF3D7_0907400), and schizont egress antigen-1, SEA1 (PF3D7_1021800) (Figure 2B). We also investigated other detectable proteins in the plasmepsin family and found that only PMIV exhibited slight stabilisation in response to WEHI-601 treatment (P<0.01), while no other plasmepsin proteins showed any stabilising effect (Table S1, Figure S2). Taken together, both the SIP and thermal-PISA experiments demonstrated that the peptidomimetic inhibitor WEHI-601 targets, and is highly selective, of PMV.

### PMV peptidomimetics do not alter the parasite metabolome

We next wanted to determine if changes in the parasite metabolome were detectable upon treatment with WEHI-601. Here we treated parasites with either WEHI-601, the inactive compound WEHI-024, or DMSO for two durations: a short 5-h treatment at 22-24 hours post invasion (hpi) or longer 16-h treatment at 6-8 hpi. In both treatment durations the overall metabolomes of WEHI-601 and control-treated parasites were unchanged, except for some short chain peptides that were significantly increased (Figure 3). In the 5-h treatment, some peptides identified to be perturbed following treatment with WEHI-601 were predicted to be derived from haemoglobin (Table S2). These peptides have previously been identified in metabolomic studies of inhibitors of haemoglobin catabolism [33]. Notably, however, most of these peptides also exhibited a significant upward trend in response to the inactive control treatment (Table S2), indicating that they are off-target and unrelated to the antiparasitic activity. In the 16-hour treatment a similar trend was observed, with the same peptides increasing in both WEHI-601 and the inactive WEHI-024 treatment. The only significantly increased peptide in WEHI-601 treatment that was not found in WEHI-024-treated samples was Ala-Cys-Pro-Ser (Table S2).

**Figure 3.**
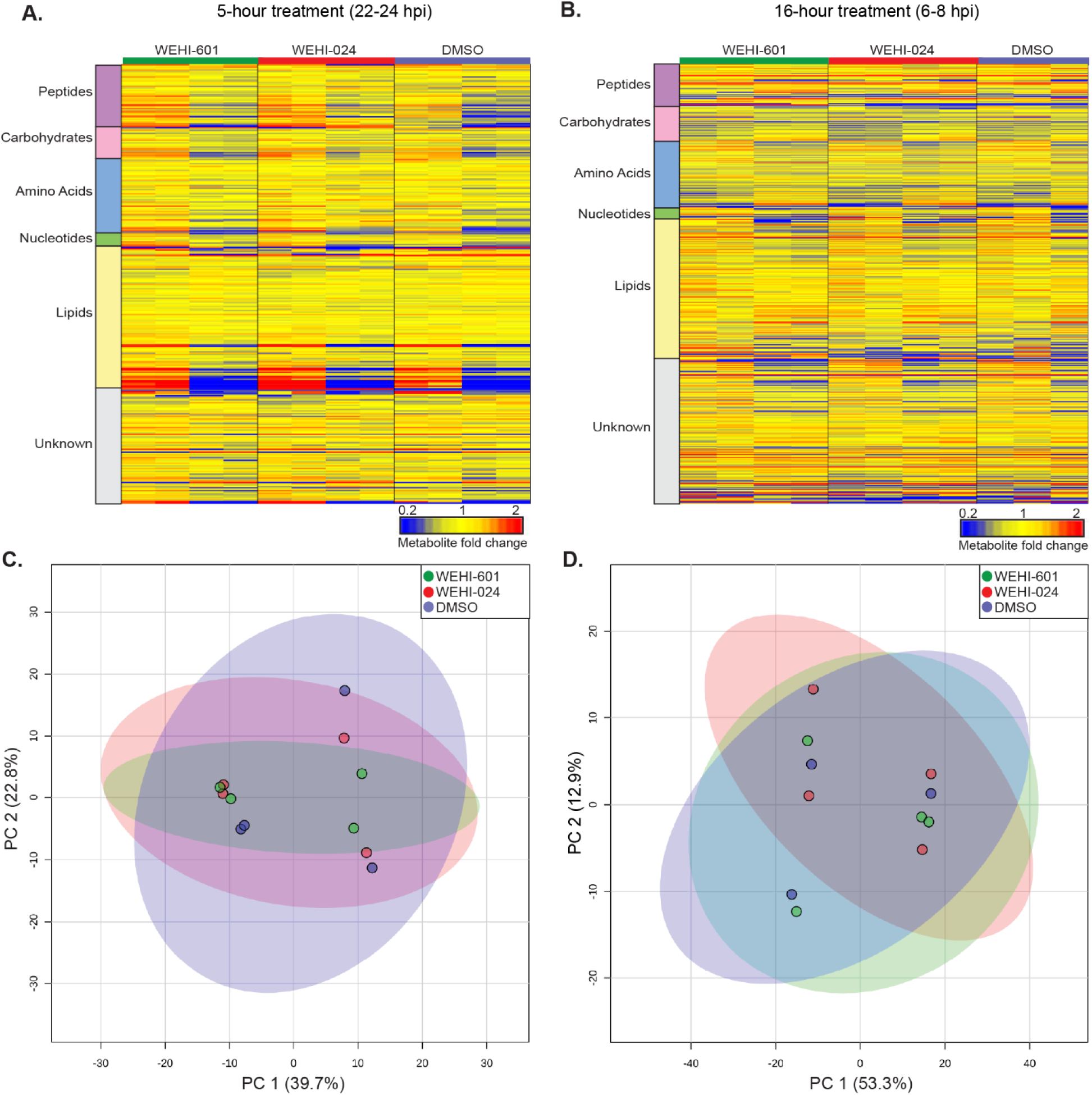
Untargeted metabolomics analysis of 3D7 parasites following treatment with WEHI- 601, WEHI-024 (negative control) or DMSO control. Heat map profile of peak intensities of all metabolites from **(A)** a 5-h drug exposure of enriched parasites at 22-24 h post invasion (hpi), or **(B)** a 16-h drug exposure of parasites at 6-8 hpi. All compounds were incubated at a concentration of 0.9 µM, heat maps show two biological replicates with two technical replicates, except for DMSO which had one technical replicate in the second biological replicate. Red, blue, and yellow indicates increase, decrease or no change in the relative abundance of metabolites based on the relative peak intensity abundance, respectively. Principle component analysis (PCA), showing scores plot for components one and two for the 5-h treatment **(C)** and 16-h treatment **(D)**. Data points indicate individual sample replicates within each treatment and the shaded area denotes 95% confidence interval.

Overall, the metabolomic study showed that the peptidomimetic PMV inhibitor WEHI-601 did not significantly affect the metabolome of *P. falciparum* 3D7 parasites compared to structurally alike PMV inactive control compound WEHI-024.

### WEHI-601 resistant parasites contain a mutation in *plasmepsin v*

To consolidate the on-target activity of the PMV peptidomimetic inhibitors, we performed a drug pressure resistance study. To generate resistance, we treated 3D7 parasites with WEHI-601 at 0.9 µM until the parasite death was observed by Giemsa-stained blood smears. The compound was then removed, and parasites were allowed to recover. This drug cycling was repeated for 3 cycles to select for resistant parasites. WEHI-601 was shown to have a 3-fold difference against the parental line (EC_50_ 0.38 µM) compared to 3D7 (EC_50_ 0.11 µM) using an LDH assay (Figure S3). Five clonal parasites strains were generated from the 3E parental strain. WEHI-601 was shown to have similar activity against each of the five clonal strains, indicating the resistance obtained was stable (Figure S3).

Three of the five resistant clones (F10, D6 and B11) were selected for whole genome sequencing. Upon subsequent re-testing of the three clonal lines, it was found that clone F10 had a greater shift in EC_50_ against WEHI-601 compared to the other two clones (Figure 4A). Analysis of the WEHI- 601 resistant genomes revealed one shared single nucleotide variant (SNV) across all three clones. This was a Thr 371 Pro mutation in the *plasmepsin v* gene (PF3D7_1323500) (Table 2). Resistant clone F10 contained additional non-synonymous SNVs in three further genes including in a *vacuolar protein sorting-associated protein 53* (PF3D7_0727000), *tRNA methyltransferase* (PF3D7_1019800) and a *cell cycle associated protein* (PF3D7_1220300) (Table 2). No additional variants across the resistant clones were detected.

**Figure 4.**
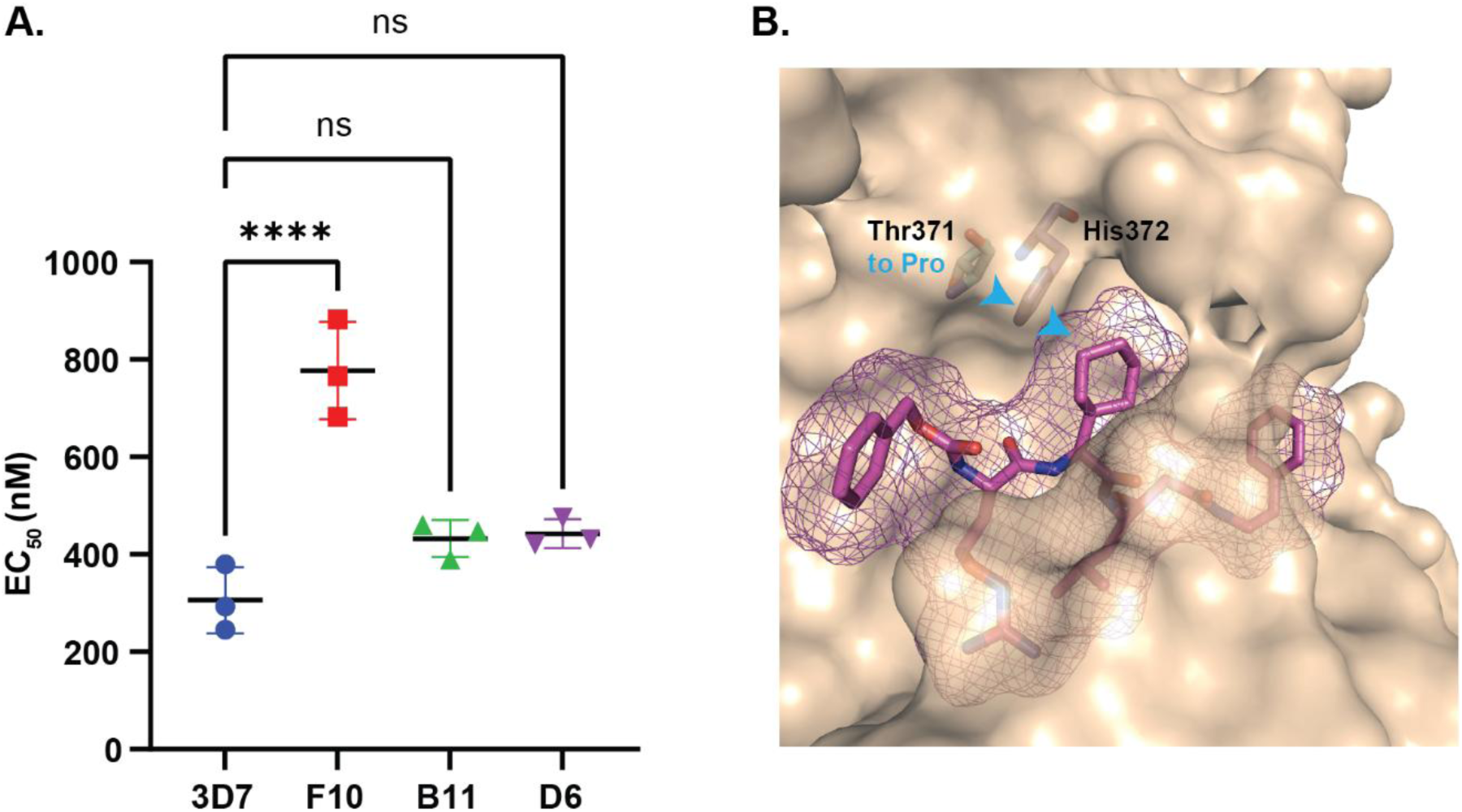
Generation of parasite resistance against WEHI-601 reveals a T371P mutation in Plasmepsin V. **(A)** Three clonal resistance lines (F10, B11 and D6) were generated against WEHI- 601 and EC_50_’s were determined in parasite growth lactate dehydrogenase assays. Error bars depict the standard deviation of 3 biological replicates. Statistical analyses via a one-way ANOVA comparing the mean of 3D7 vs. the resistant clones. **** indicates p<0.0001, ns indicates not significant. P-values for B11 and D6 compared to 3D7 were 0.11 and 0.10, respectively. Dose-response curves can be found in Figure S4A. **(B)** The structure of PvPMV in complex with WEHI- 601 demonstrating the S_2_ pocket within PMV in which the T371P mutation is located.

**Table 2.** Nonsynonymous single nucleotide variants (SNVs) found in WEHI-601 resistant clones F10, D6 and B11.^a^

| Clone | Chromosome | Position | Ref | Alt | Gene_ID | Gene_Description | Codon Change |
| --- | --- | --- | --- | --- | --- | --- | --- |
| **F10** | Pf3D7_13_v3 | 976513 | A | C | PF3D7_1323500 | plasmepsin V | T371P |
| **D6** | Pf3D7_13_v3 | 976513 | A | C | PF3D7_1323500 | plasmepsin V | T371P |
| **B11** | Pf3D7_13_v3 | 976513 | A | C | PF3D7_1323500 | plasmepsin V | T371P |
| **F10** | Pf3D7_07_v3 | 1148030 | G | A | PF3D7_0727000 | vacuolar protein sorting-associated protein 53 | P1010H |
| **F10** | Pf3D7_10_v3 | 805275 | C | A | PF3D7_1019800 | tRNA methyltransferase | E770K |
| **F10** | Pf3D7_12_v3 | 808470 | G | A | PF3D7_1220300 | cell cycle associated protein | R259K |

<sup>a</sup>Highlighted rows indicates a SNV that was shared amongst the resistant clones.

The PMV^T371P^ mutation was then mapped to the X-ray structure of *P. vivax* PMV in complex with WEHI-601 demonstrating that the mutation was localised adjacent to the S_2_ pocket of PMV which accommodates the P_2_ CyHex group of WEHI-601 (Figure 4B) [34]. This result indicates that the mutation may be induced by the unnatural CyHexGly amino acid in the P_2_ position of WEHI-601. Examining the location of this SNV across multiple *Plasmodium* spp. including *P. falciparum*, *P. vivax*, *P. knowlesi*, *P. malariae*, *P. yoelii*, *P. cynomolgi* and *P. berghei* showed that the site of the mutation and surrounding amino acids are highly conserved (Figure S4). This could indicate that this region plays an important role in PEXEL substrate specificity across *Plasmodium* species.

Since this region in PMV could be important for the enzyme activity, we determined if the mutation led to a fitness cost in parasite growth. To assess this, we grew the clones F10, B11 and D6 containing the PMV^T371P^, in addition to a wildtype 3D7 control, for three cycles of growth. Each cycle we took samples and conducted a lactate dehydrogenase (LDH) assay as a proxy for parasite growth (Figure S5A). No significant difference was observed in the amplification rate between WEHI-601 resistant clones and 3D7 (p >0.05) between cycles two and three (Figure S5B). This indicates that the T371P mutation in PMV does not induce any growth defects in parasites.

### Evaluation of PMV peptidomimetic inhibitors with varied P_2_ sidechain against WEHI-601 resistant parasites

Since no fitness cost was observed in the WEHI-601 resistant parasites, we hypothesised that the mutation would not affect the binding of natural P_2_ amino acids that are commonly found in PEXEL substrates, such as Val, to S_2_ pocket of PMV. To assess this, we tested if other PMV peptidomimetic inhibitors with smaller P_2_ side chain groups were cross-resistant to the WEHI-601 resistant parasites. These included WEHI-842 and WEHI-916, both of which have a Val in the P_2_ position; WEHI-404 with a NorVal in the P_2_ position (or n-propyl group as the P_2_ side chain) and WEHI-912 with Ile as P_2_ residue (Figure 1). To evaluate parasite growth upon treatment, *P. falciparum* 3D7 parasites were treated with these PMV peptidomimetic inhibitors in a dose response for 72 h and subsequent parasitemia was measured using an LDH assay (Figure S6). This revealed that WEHI-916, WEHI-842, WEHI-404 and WEHI-912 had equipotent activity against the WEHI-601 resistant clones F10, D6 and B11 compared to 3D7 (Table 3). This result supports that the T371P mutation in PMV effects binding of the larger unnatural CyHex group but does not significantly affect the binding of Val, NorVal or Ile, all of which contain smaller side chain groups.

**Table 3.**
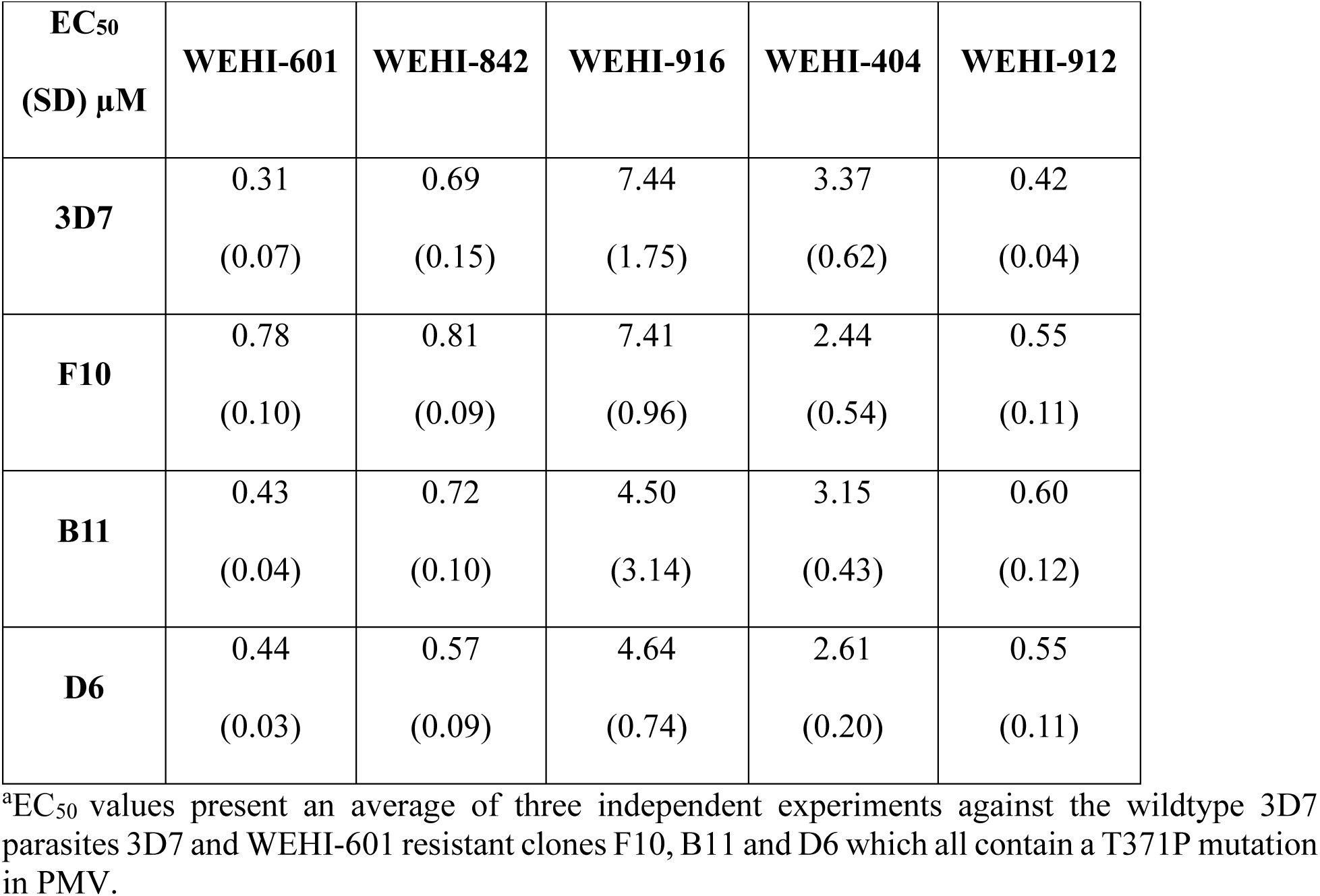
Activity of PMV peptidomimetic inhibitors with different P_2_ sidechains against WEHI- 601 resistant parasites in lactate dehydrogenase growth assays.^a^

| **EC<sub>50</sub>**<br>**(SD) μM** | **WEHI-601** | **WEHI-842** | **WEHI-916** | **WEHI-404** | **WEHI-912** |
| --- | --- | --- | --- | --- | --- |
| **3D7** | 0.31<br>(0.07) | 0.69<br>(0.15) | 7.44<br>(1.75) | 3.37<br>(0.62) | 0.42<br>(0.04) |
| **F10** | 0.78<br>(0.10) | 0.81<br>(0.09) | 7.41<br>(0.96) | 2.44<br>(0.54) | 0.55<br>(0.11) |
| **B11** | 0.43<br>(0.04) | 0.72<br>(0.10) | 4.50<br>(3.14) | 3.15<br>(0.43) | 0.60<br>(0.12) |
| **D6** | 0.44<br>(0.03) | 0.57<br>(0.09) | 4.64<br>(0.74) | 2.61<br>(0.20) | 0.55<br>(0.11) |

<sup>a</sup>EC<sub>50</sub> values present an average of three independent experiments against the wildtype 3D7 parasites 3D7 and WEHI-601 resistant clones F10, B11 and D6 which all contain a T371P mutation in PMV.

### PMV^T371P^ mediates resistance to WEHI-601

To investigate the effect of PMV^T371P^ in mediating resistance to WEHI-601, we employed reverse genetics using CRISPR-Cas9 to engineer the SNV into wildtype parasites. Here we designed a donor plasmid consisting of endogenous wildtype *plasmepsin v* homology regions (‘HR1 and HR2’) with HR1 fused to a recodonised region that contained either the wildtype PMV^T371^ or the PMV^T371P^ mutation. For expression and localisation analysis, we included a hemagglutinin (HA) tag, and a human dihydrofolate reductase expression cassette (hDHFR) to select for transfectants resistant to WR99210. A gRNA was designed binding to a protospacer adjacent motif (PAM) upstream of the SNV to mediate the Cas-9 cleavage which was then repaired through homologous recombination using donor plasmid as the template (Figure 5A).

**Figure 5.**
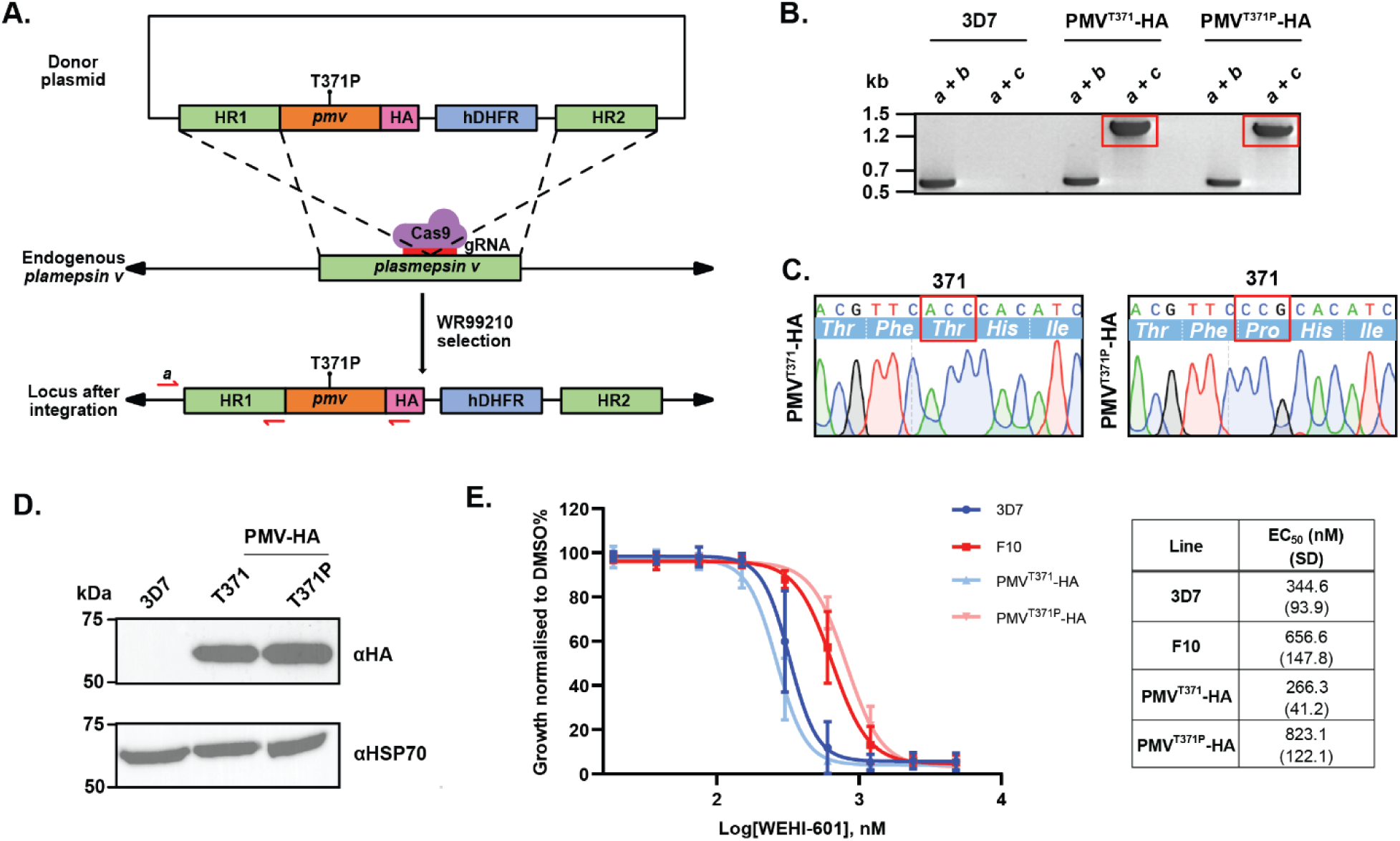
Introduction of T371P into *plasmepsin v* in 3D7 parasites mediates resistance to WEHI-601. **(A)** Design of the donor plasmid to introduce SNV (T371P) into 3D7 parasites. Homology regions (HR) were designed to be 5’ flank (HR1) and 3’ flank (HR2) where recondonised *plasmepsin v* (*pmv*) followed after HR1 (shown in orange). Cas9 was directed by a synthetic RNA to the cleavage site to perform double crossover homologous recombination. Human dihydrofolate reductase (hDHFR) was introduced to allow transfected parasites selectable by WR92210. Primers were designed to confirm correct integration, where *a* was located in the 5’ untranslated region (UTR), while *b* was located in HR1 region, and *c* was located within the haemagglutinin (HA) tag. **(B)** PCRs using these primers were performed with genomic DNA from 3D7 and both WT and mutant CRISPR lines using the primers *a, b* and *c* in (A) where the two products in the red box were sequenced. **(C)** Sanger sequencing confirmed the presence of the T371P (ACC-CCG) in PMV^T371P^-HA parasite genomic DNA. **(D)** Western blot with anti-HA demonstrated that the transfected lines contained a HA tag with the expected size of 72 kDa. Anti- HSP70 antibody was a loading control. **(E)** Dose response curves over a 72-h lactate dehydrogenase assay shows activity of WEHI-601 is reduced against both original F10 resistant clone and transfected PMV^T371P^-HA parasite lines. Error bars are SD which are indicated in brackets. EC_50_ values indicate an average value of the three independent experiments.

After transfectant parasites of both PMV^T371^-HA and PMV^T371P^-HA were successfully obtained, genomic DNA was extracted, and integration PCRs were performed using primers located in both the 5’UTR and HR1 (‘a/b’) and 5’UTR and HA tag (‘a/c’) regions (Figure 5A). This revealed that only transfected parasites with the donor plasmid could obtain a product with the PCR ‘a/c’, indicating that integration had successfully occurred (Figure 5B). These PCR products were sequenced and showed that the desired wildtype PMV^T371^ (ACC) or PMV^T371P^ (CCG) sequences were present (Figure 5C). HA-tagged protein expression was confirmed by anti-HA immunoblot (Figure 5D).

We next investigated PMV^T371^-HA and PMV^T371P^-HA parasites against WEHI-601 via a 72-h LDH growth assay. Here we included 3D7 as the parental control, in addition to the original F10 resistant clone. It was observed that there was a 3.1-fold increase in EC_50_ in the PMV^T371P^-HA parasite line when compared to the equivalent wildtype control line PMV^T371^-HA (EC_50_ 823 vs. 266 nM) (Figure 5E). The fold change in growth observed was comparable to that seen in the 3D7 wildtype line when compared to the F10 resistant clone (EC_50_ 345 vs. 657 nM) (Figure 5E). This result indicates that the T371P mutation in PMV mediates to resistance towards WEHI-601.

## DISCUSSION

Due to the prevalence of resistance to clinically used antimalarials in malaria endemic regions, the need for novel antimalarial discovery and development is urgent. PMV provides an attractive target due to its essentiality in multiple stages of the *Plasmodium* lifecycle [13, 19]. Our efforts have focussed on developing peptidomimetic compounds that mimic the natural structure of PEXEL proteins to block PMV function [27]. However, discourse has remained regarding the on-target activity of these peptidomimetic inhibitors so herein we confirmed their activity against PMV using multiple strategies.

In this study, by testing PMV peptidomimetic inhibitors WEHI-601, WEHI-842 and WEHI-916 against recombinant *Plasmodium falciparum* PMV, PMIX and PMX, we found the inhibitors presented good selectivity against PMV with low-mid nanomolar affinity. They did not display activity against PMIX but some modest affinity against PMX was observed with WEHI-842 and WEHI-916. This may be due to the structural similarity between PMV and PMX, whereby both proteins share a similar aspartic protease core structure [28]. Despite this modest activity against PMX, the selectivity index of the peptidomimetic inhibitors against PMV remains high, indicating PMV inhibition remains their main mode of action against the parasite. Intriguingly, the IC_50_ of the inhibitors against recombinant PMV did not align with the EC_50_ values against parasite growth. This could possibly be due to the inherent peptide-like characteristics of the peptidomimetics having limited cell membrane permeability resulting in modest parasite activity. Further work is required to enhance the potency of these peptidomimetic inhibitors against the parasite *in vitro*. Importantly, however, the PMV peptidomimetic inhibitors displayed little to no activity against the human aspartyl proteases, BACE1 and renin, which is advantageous for their drug development potential.

Target-compound engagement methods SIP and thermal PISA profiling both demonstrated stabilisation of PMV by WEHI-601 in both *in vitro* and live cell conditions, respectively. The second most significantly stabilised protein by WEHI-601 was SR1. In a previous study, immunoprecipitation experiments of a PMV-HA parasite line enriched SR1 [35], indicating that SR1 is likely an interactor of PMV. Since we have now shown that WEHI-601 stabilises PMV, it is probable that through its interaction with PMV, SR1 would also be stabilised in the target engagement assays. This finding underscores the value of live-cell target engagement studies in revealing physiologically relevant pathways and interactors. ClpY (PF3D7_0907400) was another protein hit that displayed significant stabilisation in the intact-cell thermal PISA profiling. ClpY is an ATPase subunit of an mitochondria ATP-dependent protease which has been shown to be important in programmed cell death in *P. falciparum* asexual stage development [36]. SEA1 or schizont egress antigen-1 (PF3D7_1021800) was also significantly stabilised by WEHI-601 which is known to act as an upstream regulator for packaging of nuclei during parasite schizogony [37]. In the absence of strong stabilisation, and since both ClpY and SEA1 are not involved in the protein trafficking pathway, these proteins are unlikely to be stabilised with direct interactions with WEHI-601.

Live cell thermal-PISA detected all 7 of the plasmepsins that are expressed in the asexual blood stage (PMI-V, IX-X). Apart from PMV, WEHI-601 failed to stabilise any other plasmepsin proteins except for some minimal stabilisation of PMIV. This may have resulted from the sequence similarity in the catalytic centre of PMV and PMIV [38]. This affinity, however, is unlikely to contribute to the mechanism of action of WEHI-601 since we have not observed any phenotype associated with haemoglobin digestion [24] and PMIV is not essential for asexual stage development [4, 6].

To complement the proteomics studies, we generated WEHI-601 resistant parasites and whole genome sequencing revealed a T371P mutation in PMV. We found this mutation site in PMV is adjacent to the H372 position which forms hydrogen bond to S368 that binds within the PfPMV catalytic aspartate D365 [38]. This implies that the region surrounding the T371P mutation site is critical to PMV function which may explain why the region surrounding the mutation is highly conserved across several different *Plasmodium* spp. We did not, however, observe any fitness cost to parasites that contained the PMV^T371P^. However, this was performed only across three cycles of growth so extending this out to evaluate if there may be a long-term growth fitness cost would be desirable. Nonetheless, it is possible that PMV^T371P^ only impairs the access of the large cyclohexane P_2_ residue of WEHI-601 to access the S_2_ pocket of PMV but the smaller natural amino acid P_2_ residues of endogenous substrate PEXEL proteins remain unaffected. Supporting this, we found no cross resistance from parasites containing the PMV^T371P^ to compounds containing P_2_ Val (WEHI-842 and WEHI-916), P_2_ NorVal (WEHI-404), and P_2_ Ile (WEHI-912).

To further investigate the role of the T371P mutation in WEHI-601 resistance, we introduced the PMV^T371P^ mutation into wildtype 3D7 parasites which demonstrated this mutation mediated resistance against WEHI-601. In the original resistant clone F10, however, we also identified three additional genes with SNVs including in the *vacuolar protein sorting-associated protein 53* gene which encodes the putative protein PfVPS53. In yeast and mammalian systems, VPS53 is a subunit of the Golgi-associated retrograde protein (GARP) complex, known to facilitate the fusion of endosome-derived transport carriers with the trans-Golgi network [39–41]. In *Plasmodium* three of the four GARP subunits are expressed [42], indicating it is likely to have a similar function in the parasite. Given that PfVPS53 may play a role in protein trafficking within the secretory pathway, it could represent a secondary mutation contributing to the mechanism of action of the peptidomimetic inhibitors. Therefore, further efforts to reverse engineer this SNV would be worthwhile. Of note, different methods for whole genome sequencing F10 and the other two clones B11 and D6 were utilised; MinIon Nanopore sequencing for clone F10; and Illumina next-generation sequencing (NextSeq) for clones B11 and D6. This could be a contributive factor to differences found in their analyses.

We also investigated whether WEHI-601 would alter the parasite metabolome through metabolomics analysis. We found no significantly distinguishable changes in metabolites between the WEHI-601 treated group, and the control treatments, indicating that the PMV peptidomimetic inhibitors do not have a significant effect on parasite metabolome. This phenomenon has been observed with other compounds that inhibit export, such as KAF156, an antimalarial agent with an unknown target in the secretory pathway, which similarly showed no effect on the parasite’s metabolome [43].

In this study, through proteomics and genetic methods, we have demonstrated the on-target activity of PMV peptidomimetic inhibitors to provide a platform for the further development of such inhibitors. However, a conditional knockdown approach has shown that parasite survival can be maintained with less than 1% of normal PMV levels [33], suggesting that PMV activity must be reduced to extremely low levels to significantly impair parasite growth. Thus, PMV peptidomimetic inhibitors require further chemical modification to improve potency and permeability before they achieve high enough efficacy to inhibit parasite growth at concentrations in line with current antimalarial compounds in the clinic. The resistant parasite lines that we selected and transfectant parasites that we created in this study can be useful in such future studies to evaluate the activity of PMV peptidomimetic inhibitors. Due to PMV’s highly conserved sequence in *Plasmodium spp.* [24] and essential function during other stages of the lifecycle [13], there is potential for PMV peptidomimetic inhibitors to evolve as a cross-species and multistage antimalarial.

## EXPERIMENTAL SECTION

### Parasite culture and lines used in this study

*P. falciparum* parasites were cultured as previously reported (Trager et al, 1976) in human O type RBCs (Australian Red Cross Blood Bank) at 4% haematocrit in RPMI-HEPES media supplemented with 5% v/v heat-inactivated human serum (Australian Red Cross Blood Bank) and 5% v/v albumax (Gibco) (herein referred to as complete RPMI) unless specified. *P. falciparum* wildtype parasites in all experiments were laboratory strain 3D7 parasites unless specified. A *P. falciparum* parasite line expressing haemagglutinin-tagged Plasmepsin X (PMX-HA) [12] was used for solvent-induced protein precipitation experiments.

### Compounds

WEHI-601, WEHI-912, WEHI-404 (referred as compound **27**, **29** and **10** in [27]), WEHI-842 [26], and WEHI-916 [24] were synthesized as reported before. The above compounds were all dissolved in DMSO to a 10 mM stock solution and stored at -20°C.

### Plasmepsin biochemical assays

This was performed as per [44] with the following changes. Starting concentrations of compounds were as follows: 10-100 μM (PvPMV assays), 1-11 μM (renin, cathepsin D and BACE-1 assays) or 100 μM (PfPMX and PfPMIX assays).

### Lactate dehydrogenase (LDH) Malstat growth assay

These assays were performed as previously described [45]. Synchronised ring-staged parasites were treated with serial diluted compounds at 0.5 % parasitemia with 2% haematocrit in 100 µL final volume in 96-well plates. A nine-point titration was performed with a 1 in 2 serial dilution in complete RPMI media with starting concentrations of WEHI-601 (4.8 µM), WEHI-912 (5 µM), WEHI-404 (20 µM), WEHI-842 (20 µM), and WEHI-916 (20 µM). Plates were incubated at 37°C for 72 h before parasitemia was quantified using a lactate dehydrogenase (LDH) malstat assay as previously described [27]. Dose-response curves and EC_50_s were analysed through GraphPad Prism 10.3.0 using four-parameter log(inhibitor) vs. response nonlinear regressions.

### WEHI-601 resistant parasite generation

This was performed as previously described [46] with a selection pressure of 0.9 µM which equated to approximately 10 x EC_50_.

### Fitness cost parasite growth assay

WEHI-601 resistant clones F10, B11 and D6, and a 3D7 parasite line were synchronised via 5% sorbitol lysis 48 h before assay set up. For assay set-up, ring-stage parasites were adjusted to 0.5% parasitemia and 2% haemotocrit and incubated at 37°C. Samples were then collected each cycle for 3 cycles and stored frozen until assay completion. To prevent parasite overgrowth, cultures were diluted 1:8 at each cycle, with this dilution factored into the analysis. Parasite growth was quantified using the LDH assay described above.

### Whole genome sequencing of clone F10 WEHI-601 resistant parasites

This was performed as previously described [47] with alterations outlined below. The PMV isolate and 3D7 control isolate were sequenced in one sequencing run and performed on a MinION platform with MIN106/R9.4.1 flow cells and MinIT (Software 18.09.1) to generate fast5 files that were then basecalled using (Guppy V3.1.5+781ed57) and demultiplexed using Porechop (V0.2.4).

Reads were aligned against the *P. falciparum* 3D7 reference genome (PlasmoDB version 31) using minimap2 (V2.1.7) using the map-ont preset. Candidate SNVs in subtelomeric, centromeric, or hypervariable regions were removed using bedtools subtract (V2.26.0) to only retain SNVs in the core genome [48]. To examine structural variants, demultiplexed fastq files were additionally aligned using a sensitive long read aligner NGMLR V0.2.7 [49]; (https://github.com/philres/ngmlr) and again processed with samtools utilities (V1.7) to sort and index the alignment files. The aligned files were then examined for structural variants using Sniffles V1.0.11 ([49] https://github.com/fritzsedlazeck/Sniffles) and filtered to contain high read support >=10 and filtered based on structural variant length >=200bp. SNVs in 12 genes that were common to both the F10 WEHI-601 resistant genome and an unrelated resistant genome sequenced concurrently were excluded from the analysis. The data for this study have been deposited in the European Nucleotide Archive (ENA) at EMBL-EBI under accession number PRJEB82704.

### Whole genome sequencing of clones D6 and B11 WEHI-601 resistant parasites

This was performed as previously described [50] with the following changes: Copy number analysis was performed using the R package QDNaseq v1.28.0 [51] with 1 kbp windows. Copy number windows were filtered to exclude regions that were centromeric, telomeric, or had mappability of less than 0.5 based on 30mers generated by GenMap [52]. The data for this study have been deposited in the European Nucleotide Archive (ENA) at EMBL-EBI under accession number PRJEB82800.

### Genetically engineered T371P PMV mutant *P. falciparum* parasites

To introduce the WEHI-601 resistant mutation into 3D7 wildtype parasites, 533 bp of endogenous *plasmepsin v* (PF3D7_1323500) was synthesised as homology region 1 (‘HR1’) along with the recodonised following endogenous sequence including either the WT or T371P mutation with a 3x haemagglutinin tag at the 3’ end. The entire ‘HR1-*pmv*-HA’ fragment was synthesised (GenScript) and Gibson assembly (Gibson Assembly Master Mix, New England Biolabs) was performed to insert this fragment into parasite vector p1.2 [35]. Primers for Gibson assembly were designed through NEBuilder online program https://nebuilder.neb.com/#!/ (listed in Table S3).

The guide RNA sequences (stated in Table S3) were designed through online program https://chopchop.cbu.uib.no with a protospacer adjacent motif (PAM) (Table S3). The gDNA fragment was fused into pUF1-Cas9G plasmid [35] which included the Cas-9 gene (In-Fusion HD cloning kit, Takara).

The donor plasmid was linearised and transfected with the guide plasmid into late schizont-stage 3D7 parasites (Amaxa Basic Parasite Nucleofector Kit 2, Lonza) as described in [35]. Parasites that had integrated the donor plasmid containing a human dihydrofolate resistance (hDHFR) gene into their chromosome were selected for with 2.5 nM WR99210. Once viable parasites were obtained, genomic DNA was extracted, and integration was confirmed via PCR using primers listed in Table S3. These PCR products were then Sanger sequenced (Australian Genome Research Facility) to confirm the presence of the desired mutation or wildtype sequences.

### Solvent-induced precipitation protein profiling of plasmepsin V

Lysate of the *Pf*PMX-HA parasite line was prepared as described in [53] without the addition of protease inhibitors. Briefly, parasite lysate was treated either with WEHI-601 (10 µM) or DMSO (0.1% v/v) for 3 min before being aliquoted into 0-25% acetone/ethanol/formic acid mixture (AEF) at a 50:50:1 (v/v) solution. After agitation at 800 rpm for 20 mins at 37°C, the precipitated protein was removed by centrifuge at 17 000 g for 20 min at 4°C and supernatant was transferred into 1x NuPAGE LDS Sample Buffer (Invitrogen) with 1:100 2-mercaptoethanol (Sigma Aldrich). The sample was boiled for 3 min and proteins were separated on 4-12% acrylamide gels (NuPAGE, Invitrogen). Blots were then probed with anti-PMV antibody (1:1000) [24], anti-HA (1:1000) (Roche) and anti-*Pf*START1 (1:1000) [53] with corresponding HRP antibodies (1:2000) (anti-rabbit-HRP, Merck; anti-rat-HRP, Roche).

### Intact-cell thermal PISA profiling with WEHI-601

#### DIA Thermal PISA (Proteome Integral Solubility Alteration) assay

The experiment was carried out in 4 biological replicates. Synchronised mature-stage parasites (35-41 hpi) were exposed to WEHI-601 or the DMSO [Sigma] vehicle control for 1 h at standard culture conditions. Parasites were pelleted through centrifugation and resuspended in WEHI- 601/DMSO-supplemented PBS, each split into 13 identical aliquots and transferred onto a 96-well plate. Samples were heated in a PCR machine [Biorad] for 3,5min to different temperatures across 50-72°C gradient (at 2°C intervals) or to 37°C as a non-denaturing control. Cells were lysed by 3x flash freeze/thawing using liquid N_2_, followed by 10x mechanical sheering with a 29-gauge needle-syringe, and denatured protein removal through filtration at 0.2 µM level. The soluble phase was recovered and pulled together in equivolume ratios into two samples; Gradient 1: 50- 60°C and Gradient 2: 62-72°C, respectively. Sample preparation for proteomic analysis was carried out using modified SP4 protocol [54]. In brief, 20 µg of protein was reduced [20 mM TCEP, 100 mM TEAB] for 10 min at 95°C and alkylated with 55 mM CAA for 30 min. Following addition of 20 µL of PureCube Carboxy magnetic beads [Cube biotech] and neat ice-cold ACN to a final concentration of 80%, samples were incubated on a thermomixer for 20 min at RT at 800 RPM and pelleted down at 3000g for 5 min. Beads were washed 3x with 80% Ethanol and following SN removal dried in a SpeedVac. Dried beads were subjected to sequential digestion with LysC (3h, 1:50 enzyme:protein ratio) and trypsin (overnight, 1:50 enzyme:protein ratio). Resulting digest was acidified with TFA to final 1% concentration and desalted on T3 C18 stage tips [Affinisep] according to manufacturer’s instructions.

#### MS data acquisition and data analysis

Following resuspension in 0.1% Formic Acid, 2% ACN, peptide samples were loaded on to a C18 fused silica column (inner diameter 75uM, OD360 x 15cm length, 1.6uM C18 beads) packed into emitter tip [IonOptics] and separated on a 45 min analytical gradient on a Neo Vanquish liquid chromatography system [Thermo Scientific] interfaced with MS [Orbitrap Eclipse Tribrid Mass Spectrometer, Thermo Scientific] and analysed in a DIA mode. Peptide identification was carried out in DIA-NN 1.8.1 using is silico spectral library generated from Uniprot *P. falciparum* (UP000001450) and human (UP000005640) reference proteomes. Two missed cleavages and 2 variable modifications [ox(M) and Ac(N-term)] were allowed. Differential abundance data analysis (moderated t-test) [55] of *P. falciparum* proteins was conducted in the R environment using precursor normalised MaxLFQ data for proteins detected with ³ 2 peptides. Hit selection criteria included p value <0.01, >0.2 log2 fold change in protein abundance and protein detection across all samples in the comparison.

### Metabolomics

#### Cell culture and drug incubations for LC-MS metabolomics analysis

3D7 parasites were grown in RPMI medium containing hypoxanthine and 0.5% (wt/vol) Albumax (Gibco). For the 5-h treatment, parasites were synchronised by double 5% (wt/vol) sorbitol lysis, 14 h apart and cultured for a further 52 h to ensure the experiment was performed on early-trophozoite stage cultures ∼22-24 hpi. Parasites were adjusted to 6% parasitaemia and treated with WEHI-601 or WEHI-024 (0.9 µM) for 5 h. Following the treatment duration, parasites were harvested using magnet separation to achieve enriched trophozoite samples (>80% parasitaemia). For the 16-h treatment, parasites were tightly synchronised to 0 ± 2 h by magnet harvesting late stage segmented schizonts, adding the magnet elute to uninfected red blood cells (RBCs) and performing sorbitol lysis 2 h later. Parasites were cultured for a further 6 h (∼6-8 hpi) before treatment with WEHI-601 or WEHI-024 (0.9 µM) for 16 h. Both experiments included untreated controls that contained equivalent amounts of dimethyl sulfoxide (DMSO; as a vehicle).

#### Sample extraction and LC-MS metabolomics analysis

After drug treatments, metabolites of infected RBCs (iRBCs) were collected as previously described [56], with minor modifications. Metabolites of iRBCs in the 5 h and 16 h treatments were extracted in either 100 µl or 200 µl of methanol, respectively.

Metabolite analysis was performed by liquid chromatography-mass spectrometry LC-MS using hydrophilic interaction liquid chromatography (HILIC) and high-resolution (Q-Exactive Orbitrap, Thermo Fisher) MS as previously described [56, 57]. Briefly, samples (10 μL) were injected onto a Dionex RSLC U3000 LC system (Thermo) fitted with a ZIC-pHILIC column (5 μm particle size, 4.6 by 150 mm; Merck) and 20 mM ammonium carbonate (A) and acetonitrile (B) were used as the mobile phases. A 30 min gradient starting from 80% B to 50% B over 15 mins, followed by washing at 5% B for 3 mins and re-equilibration at 80% B, was used. MS with a heated electrospray source operating in positive and negative modes (rapid switching) and a mass resolution of 35,000 from m/z 85 to 1275. Sample injections within the experiment were randomized to avoid any impact of systematic instrument drift on metabolite signals. Retention times for ∼350 authentic standards were checked manually to aid metabolite identification.

#### LC-MS metabolomics data processing

Metabolomics data sets were analysed using IDEOM [58]. Raw files were converted to mzXML with msconvert [59], extraction of LC-MS peak signals was conducted with the Centwave algorithm in XCMS, alignment of samples and filtering of artifacts with mzMatch, and additional data filtering and metabolite identification was performed in IDEOM (supplementary Data file S1). In the 5 h and 16 h-treatments, 411 and 453 putative metabolites were identified, respectively. Metabolite abundance was determined by LC-MS peak height and was normalized to the average for untreated samples. Statistical analyses used Welch’s t test (p-value< 0.05) and Pearson’s correlation (Microsoft Excel). Principal-component analysis (PCA) was generated in Metaboanalyst, a web interface [60].

## ASSOCIATED CONTENT

### Supporting Information

Supporting Information files contain dose-response curves of selected compounds against *P. falciparum* parasites, and other supplementary content. “This material is available free of charge via the Internet at http://pubs.acs.org.”

## AUTHOR INFORMATION

Corresponding Authors Dr. Madeline G. Dans.

Dr. Brad E. Sleebs. Phone: +61 3 9345 2718.

## NOTES

The authors declare no conflict of interest with this manuscript.

## ACKNOWLEDGMENTS

This work was funded by the National Health and Medical Research Council of Australia (Development Grant 2014427 to B.E.S. and A.F.C.), the Victorian State Government Operational Infrastructure Support and the Australian Government NHMRC IRIISS. We thank and acknowledge the Australian Red Cross Lifeblood for the provision of fresh red blood cells, without which this research could not have been performed. We thank the University of Melbourne for provision of a Research Scholarship to W.S. A.F.C. is a Howard Hughes International Scholar and an Australia Fellow of the NHMRC. B.E.S. is a Corin Centenary Fellow.

