## Supplementary file for "Deconvolution of the on-target activity of plasmepsin V peptidomimetics in *P. falciparum* parasites"

**Index**


| Page | | |
|---|---|---|
| S3 | Figure S1 | Replicates blots of SIP assays |
| S4 | Figure S2 | Plasmepsins in thermal proteome profiling assay |
| S5 | Figure S3 | Clonal resistance to WEHI-601 |
| S6 | Figure S4 | *Plasmodium* species sequence alignment of the PMV T371 region |
| S7 | Figure S5 | Fitness cost of PMV(T371P) in parasites |
| S8 | Figure S6 | Dose-response curves of PMV(T371P) parasites against $P_2$ residues |
| S9 | Figure S7 | Uncropped blots used in this study |
| S12 | Table S2 | Untargeted metabolomics analysis of peptides |
| S13 | Table S3 | Sequences and primers used to construct donor plasmids |

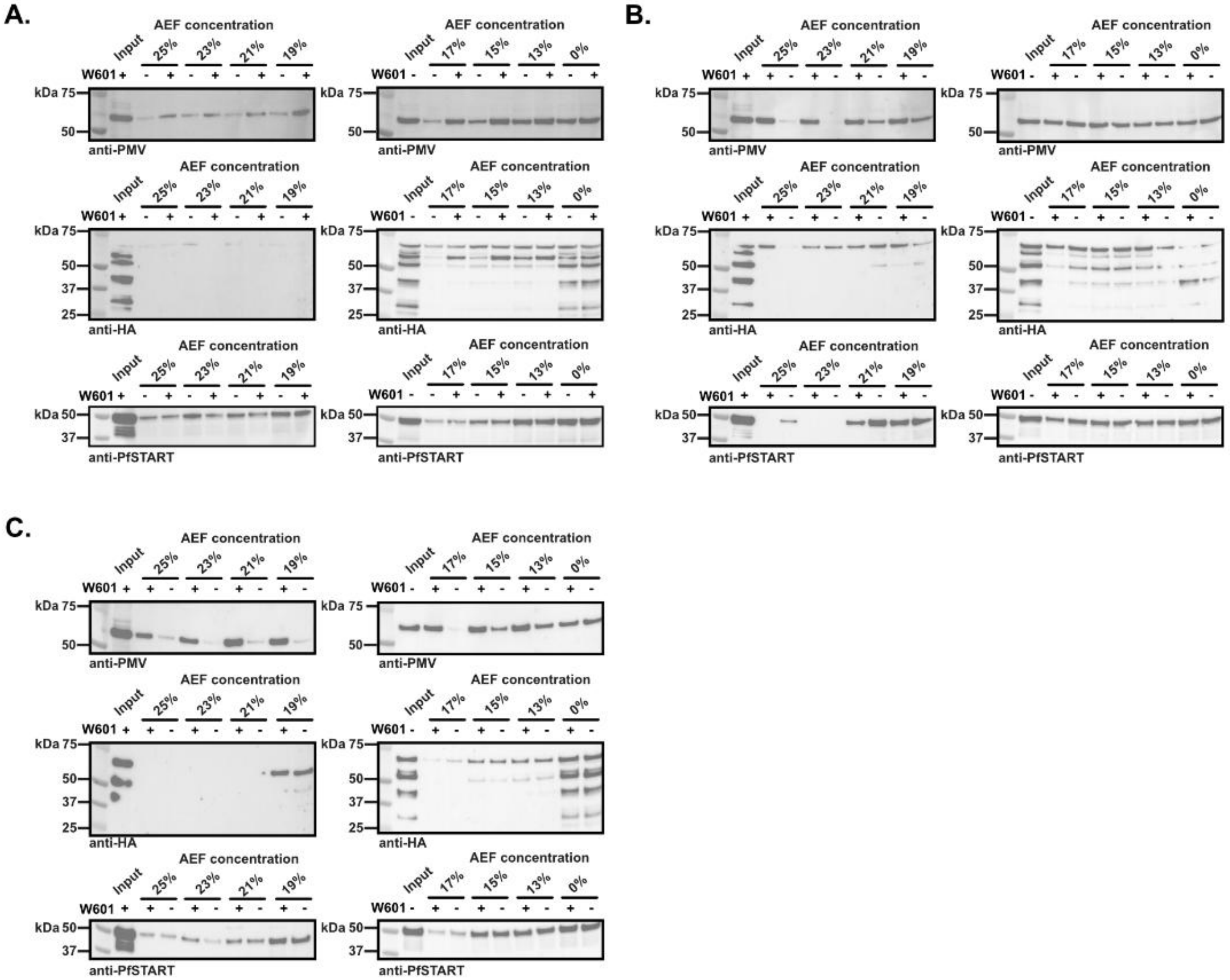


**Figure S1.** Replicates of SIP assays. The parasite lysate was treated with DMSO ('-') or 5 µM WEHI-601 ('+'). The lysate was challenged in acetone/ethanol/formic acid mixture (AEF, v:v:v = 50:50:1) in a 0-25% gradient. The soluble fraction of protein was extracted and soluble proteins were separated out via western blot and probed with an anti-PMV or an anti-HA (for PMX detection) antibodies. An anti-*Pf*START1 antibody was used as a loading control.

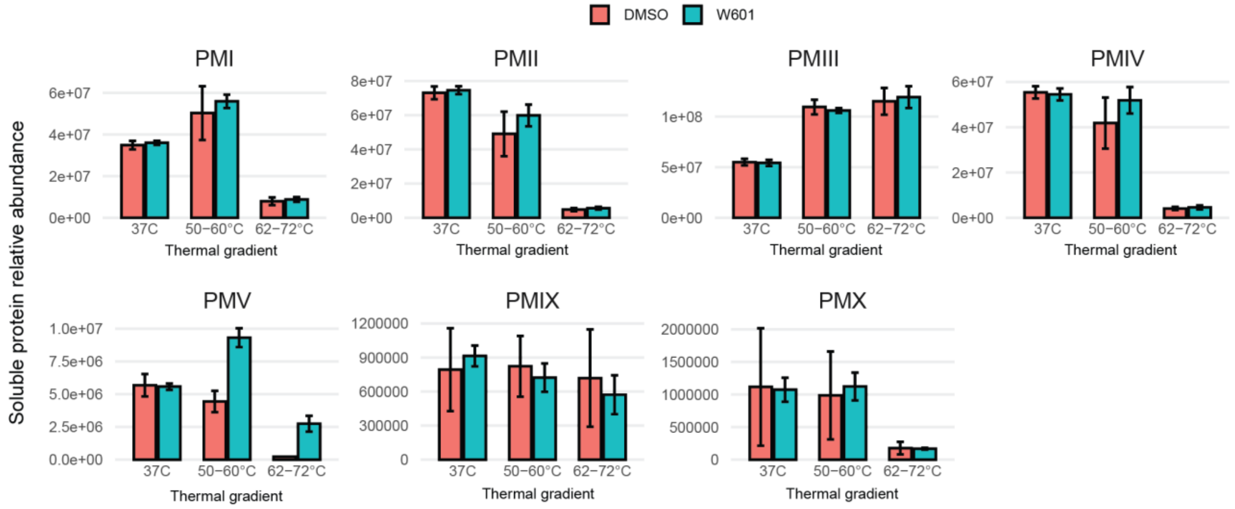


**Figure S2.** Soluble protein relative abundance of detectable *P. falciparum* plasmepsin proteins in thermal stability assays. Schizont-staged parasites were cultured with WEHI-601 (10 µM) or the DMSO control and were lysed and challenged via a thermal temperature gradient (n = 4 biological replicates, mean +/− SD). PMV and PMIV were the only plasmepsins to show significant stabilisation ($p$<0.01, actual values can be found in Table S1). Bar graphs represent the average value of 4 biological replicates.

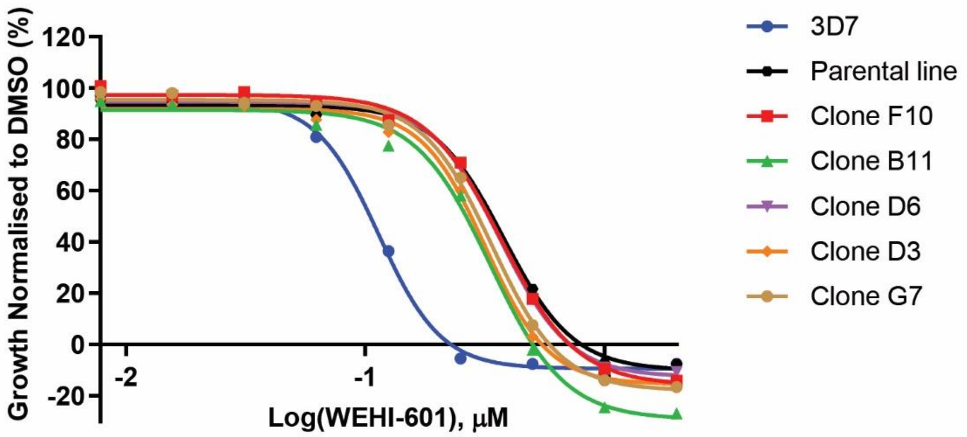


| | 3D7 | Parental line | Clone F10 | Clone B11 | Clone D6 | Clone D3 | Clone G7 |
|---|---|---|---|---|---|---|---|
| $EC_{50}$ (μM) | 0.11 | 0.38 | 0.37 | 0.33 | 0.37 | 0.32 | 0.34 |

**Figure S3.** Resistance against WEHI-601 remained stable after clonal parasites were obtained. 72-h lactate dehydrogenase growth assays determined $EC_{50}$ values of WEHI-601 against each clonal line, the parental resistant line and 3D7 wildtype parasites. Each data point represents the average of >2 technical replicates. Biological replicates of the lines selected for whole genome sequencing are represented in Figure 4 and Table 3.

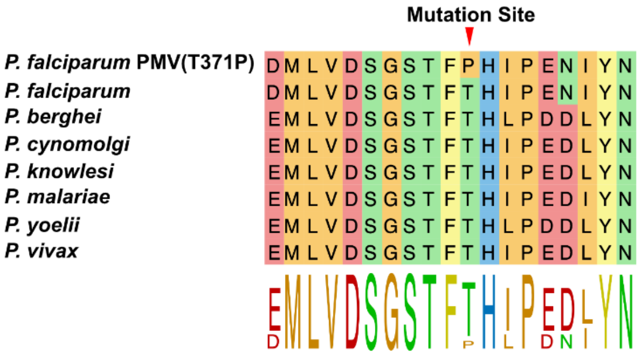


**Figure S4.** Alignment of the Plasmepsin V amino acid sequence around T371P mutation site in multiple *Plasmodium* species including *P. falciparum*, *P. berghei, P. cynomolgi, P. knowlesi, P. malariae, P. yoelii, and P. vivax.* Amino acids surrounding the T371P SNV are highly conserved. Wildtype *plasmepsin v* gene sequences were derived from PlasmoDB (rel. 9.0, 2012-MAY-18) and the alignment was performed through DNASTAR MegAlign Pro.

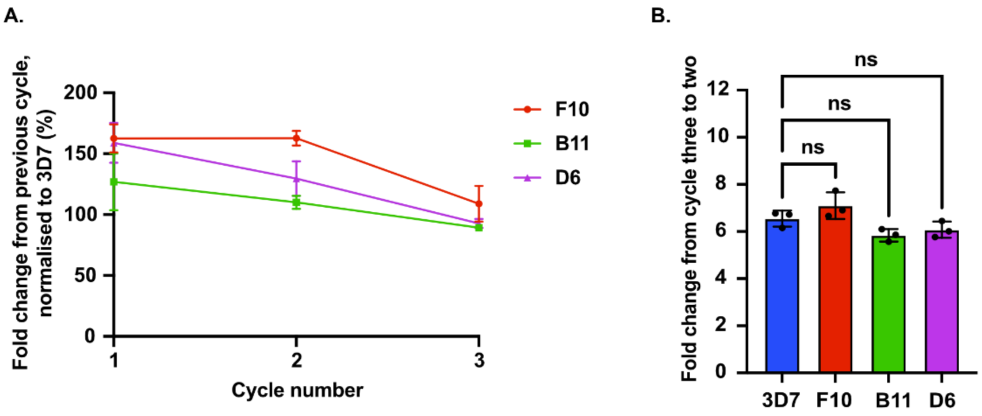


**Figure S5. (A)** Comparison of the replication rate between wildtype 3D7 and WEHI-601 resistant parasite clones containing $PMV^{T371P}$. Each cycle for three cycles, parasitemia was measured by a lactate dehydrogenase assay. **(B)** The fold change of amplification rate between cycles two and three revealed there was no significant difference ($p > 0.05$) between the resistant clones and 3D7 via a one-way ANOVA analysis. Each data point represents an average of three technical replicates. Error bars indicate the standard deviation of three technical replicates.

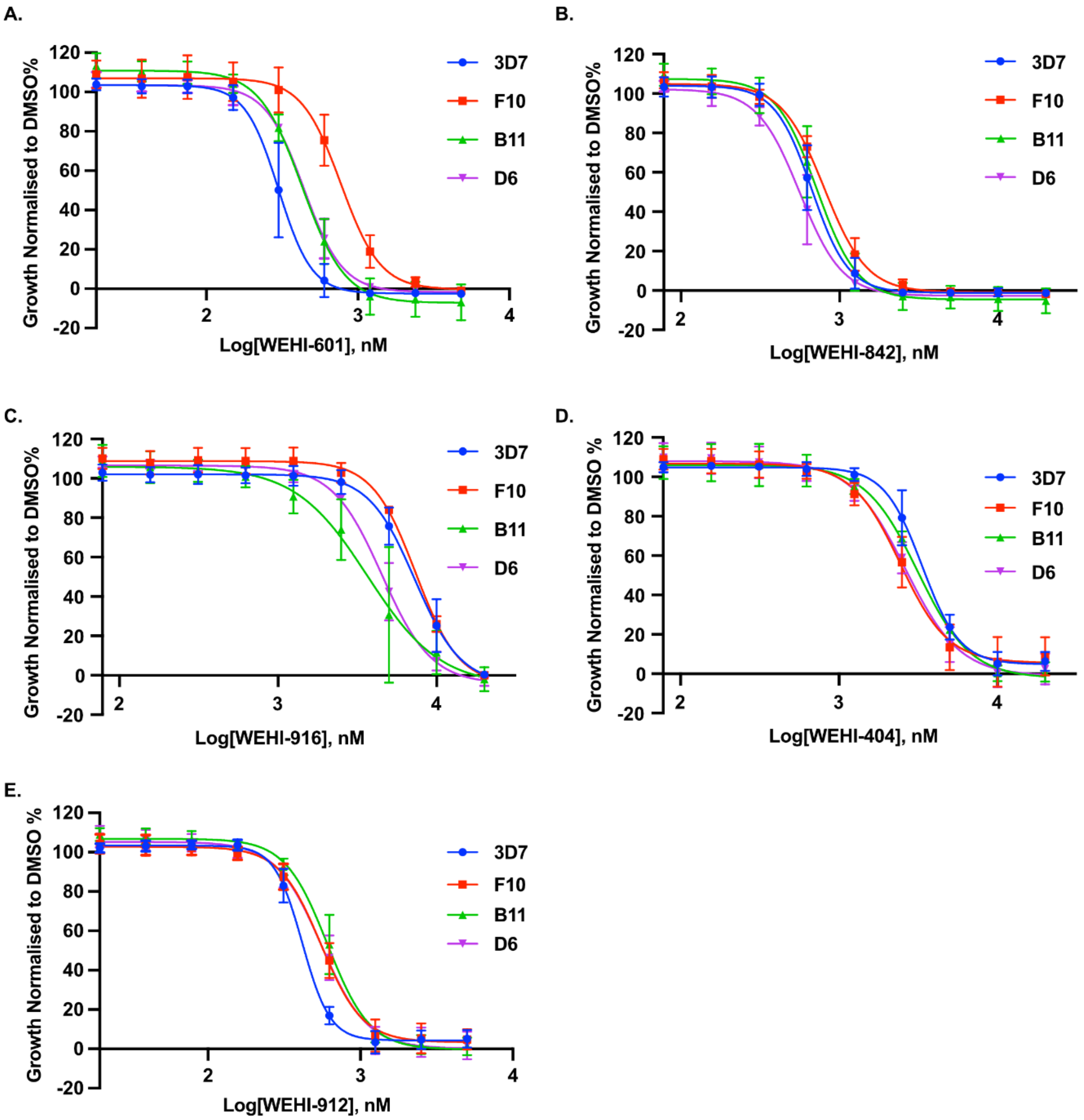


**Figure S6.** Dose response curves against WEHI-601 resistant parasite populations of **(A)** WEHI-601 with $P_2$ cyclo-hexane sidechain, **(B)** WEHI-842 with valine $P_2$ residue, **(C)** WEHI-916 with valine $P_2$ residue, **(D)** WEHI-404 with n-propyl $P_2$ side chain, and **(E)** WEHI-912 with isoleucine $P_2$ residue. F10, B11 and D6 are WEHI-601 resistant parasites comprising a PMV(T371P) mutation. $EC_{50}$ values represent an average of 3 biological replicates generated from a growth lactate dehydrogenase assays. Dose-response curves were generated in GraphPad Prism 10 using nonlinear regression curves. Error bars represent the standard deviation (SD). $EC_{50}$ and SD values can be found in Table 3.

### First replicate of SIPP

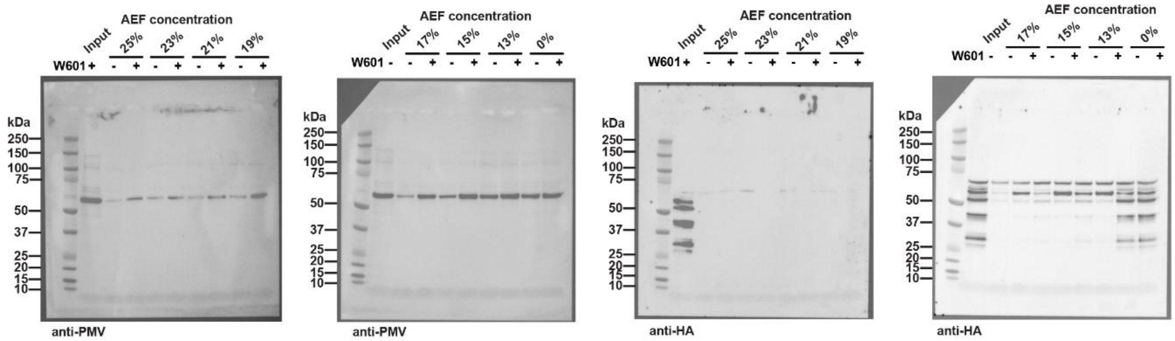


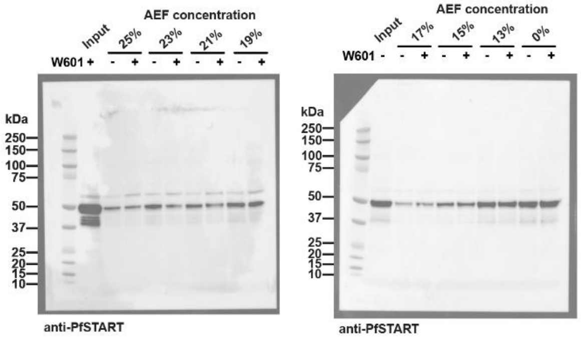


**Blots are used in Figure S2**

### Second replicate of SIPP

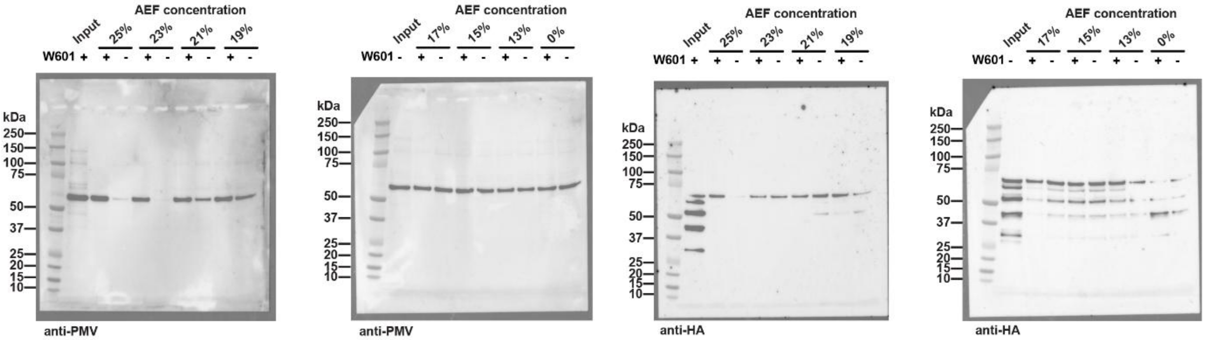


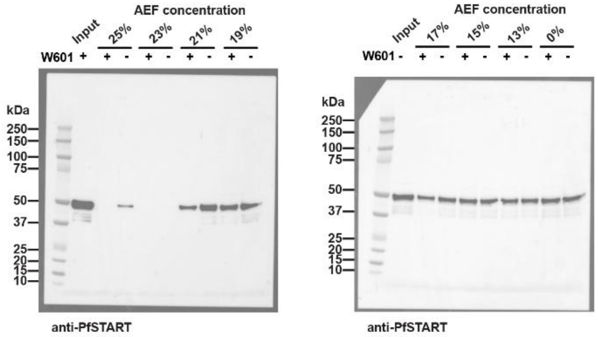


**Blots are used in Figure S2**

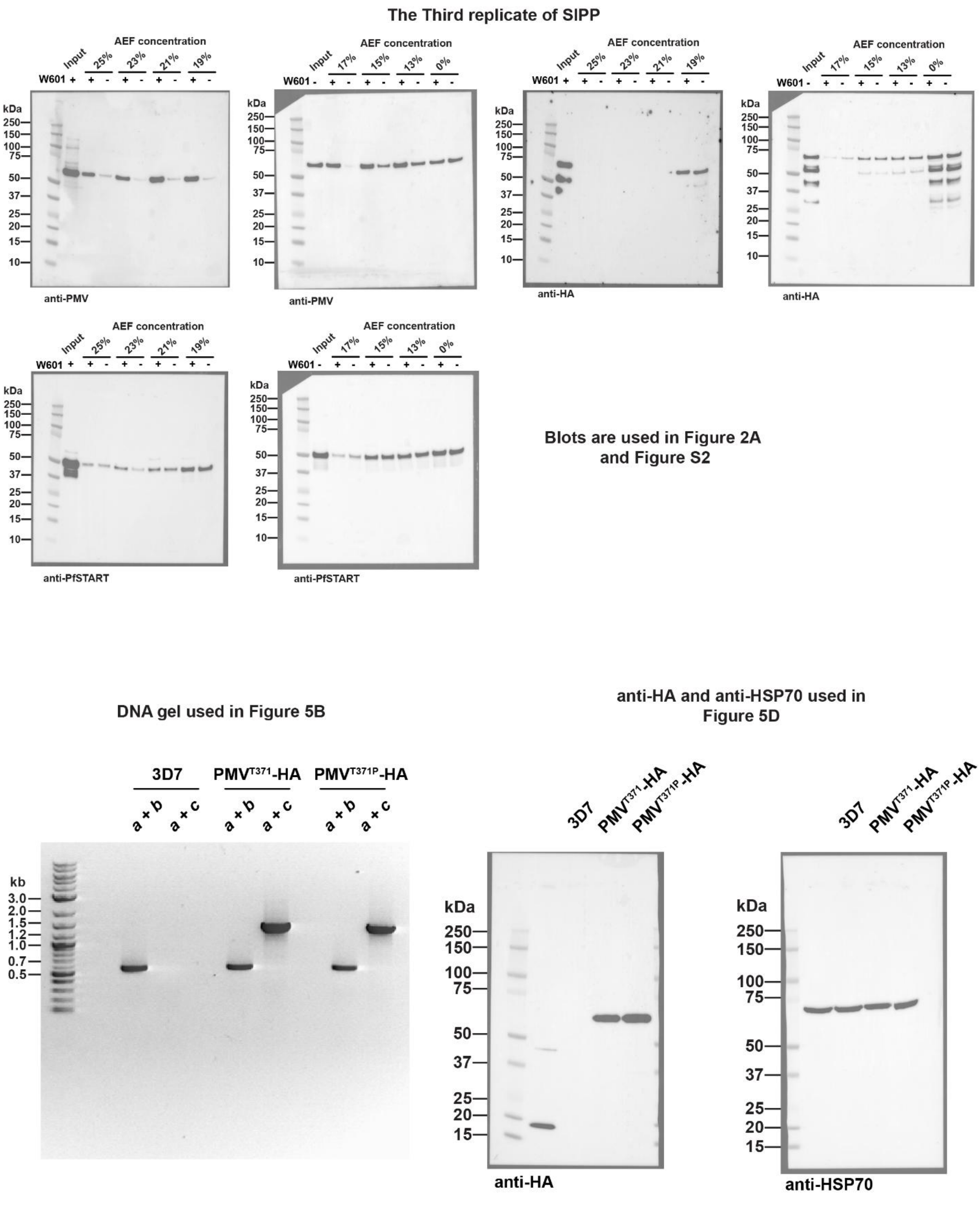


**Figure S7.** Uncropped blots used in this study.

### SUPPLEMENTARY TABLES

**Table S1. Complete hit list from thermal stability assays with WEHI-601.**

(Separate excel document)

**Table S2.** Untargeted metabolomics analysis of peptides originating from 3D7 parasites after exposure to WEHI-601 or WEHI-024. Heat map shows all statistically significant peptide abundances in mean fold change (compared to DMSO control) from a 5-h treatment at 22-24 hours post invasion (hpi) and a 16-h treatment at 6-8 hpi.[a]

| | 5-h treatment (22-24 hpi) | | | | | 16-h treatment (6-8 hpi) | | | | |
|---|---|---|---|---|---|---|---|---|---|---|
| **Metabolite Name** | **Neutral mass** | **Retention time (min)** | **WEHI-601** | **WEHI-024** | **DMSO** | **Neutral mass** | **Retention time (min)** | **WEHI-601** | **WEHI-024** | **DMSO** |
| **Dipeptides** | | | | | | | | | | |
| Ala-Ser | 176.0797 | 13.34 | **1.16** | **1.29** | **1.00** | 176.0797 | 13.34 | 1.07 | 0.93 | 1.00 |
| *Ala-Pro | 186.1006 | 11.37 | 1.01 | 1.27 | **1.00** | 186.1005 | 13.57 | 1.53 | 0.75 | 1.00 |
| Ile-Ala | | | | | | 202.1318 | 11.66 | **1.67** | **1.44** | 1.00 |
| Ile-Thr | 232.1422 | 7.59 | **2.38** | **1.94** | **1.00** | | | | | |
| Ile-Asn | 245.1375 | 9.57 | **1.38** | 1.20 | **1.00** | | | | | |
| *Lys-Val | 245.1738 | 13.43 | **1.87** | **1.86** | **1.00** | 245.1739 | 21.54 | 1.01 | 0.93 | 1.00 |
| *Glu-Thr | 248.1008 | 14.60 | 1.05 | 1.17 | **1.00** | 248.1008 | 14.54 | 1.00 | 0.91 | 1.00 |
| *Pro-His | 252.1223 | 12.19 | **2.19** | 1.55 | **1.00** | | | | | |
| *His-Leu | 268.1533 | 7.85 | **1.58** | **1.55** | **1.00** | | | | | |
| **Tripeptides** | | | | | | | | | | |
| Ala-Gly-Ser | 233.1012 | 12.78 | **2.27** | **2.03** | **1.00** | | | | | |
| Glu-Ala-Val | 317.1586 | 10.96 | 1.22 | **1.32** | **1.00** | | | | | |
| **Tetrapeptides** | | | | | | | | | | |
| Ala-Ala-Ala-Pro | 328.1743 | 9.95 | **8.94** | 4.34 | **1.00** | | | | | |
| Ala-Ala-Ala-Ser | | | | | | 318.1539 | 15.98 | **0.73** | **0.63** | 1.00 |
| Ala-Val-Gly-Pro | 342.1899 | 9.46 | 1.23 | **1.41** | **1.00** | 342.1904 | 9.42 | 0.87 | 0.77 | 1.00 |
| Ala-Val-Gly-Pro | 342.1899 | 9.46 | 1.23 | **1.41** | **1.00** | | | | | |
| Ala-Asp-Gly-Pro | 358.1492 | 15.72 | **7.67** | 3.43 | **1.00** | 358.1491 | 15.61 | **50.74** | **2.80** | **0.00** |
| Ala-Cys-Pro-Ser | | | | | | 376.1416 | 14.57 | **3.25** | **0.00** | **0.00** |
| *Asn-Leu-Pro-Pro | 439.2430 | 10.30 | **1.31** | **1.45** | **1.00** | 439.2431 | 10.25 | 0.99 | 0.85 | 1.00 |
| Ala-Glu-Glu-His | 484.1920 | 15.79 | **1.70** | **1.46** | **1.00** | 484.1921 | 15.70 | 1.07 | 0.90 | 1.00 |
| Gln-Leu-Lys-Tyr | 530.2485 | 8.15 | **2.38** | 1.98 | **1.00** | 550.3115 | 17.42 | 1.16 | 0.75 | 1.00 |
| *Ala-Leu-Asn-Pro | 413.2272 | 9.04 | **1.53** | **1.45** | **1.00** | | | | | |
| *Ala-Leu-Trp-Ser | 475.2433 | 7.60 | **0.63** | 1.13 | **1.00** | | | | | |
| **Pentapeptides** | | | | | | | | | | |
| His/Ile/Ile/Arg/Arg | 693.4396 | 10.34 | **0.85** | 0.90 | 1.00 | | | | | |
| Ile/Ile/Val/Trp/Tyr | 709.4152 | 4.12 | **2007.18** | **245.51** | **0.00** | | | | | |

[a] Red, blue and yellow indicates increase, decrease or no change in abundance respectively. **Bold** = significant, p-value< 0.05. *denotes possible hemoglobin-derived peptide.

**Table S3. Sequences and primers used to construct donor plasmids.** Text in lower case indicates restriction enzyme cutting sites. Text italicised indicates the protospacer adjacent motif (PAM) site. Text underlined indicates the homologous arm for In-Fusion reaction. Text in matched colour indicates the homologous ends for Gibson assembly.

| | | | |
|---|---|---|---|
| **Primers to introduce synthetic PMV gene into p1.2 vector** | | | |
| Synthetic HR1-Recodon block | Insert forward | GTGACACTATAGAATACTCAAGCTGCGGCCGCTCATCTATTTTATATTG | |
| | Insert reverse | TGCCATATCCCTCGACCCGGGTTACTATCATTAGGCATAATCTGG | |
| Vector part 1 | Vector P1 forward | TTATGCCTAATGATAGTAACCCGGGTCGAGGGATATGG | |
| | Amp reverse | GTGCAAAAAAGCGGTTAGCTCCTTCGGTCCTCCGATCGTTGTCAGAAGTAAGTTG | |
| Vector part 2 | Amp forward | CGAAGGAGCTAACCGCTTTTTTGCACAACATGGGGGATCATGTAACTCGCCTTGAT | |
| | Vector P2 reverse | ATAAAATAGATGAGCGGCCGCAGCTTGAGTATTCTATAGTGTCACCTAAATAGCTTG | |
| **Primers to amplify HR2 from gDNA** | | | |
| HR2 | PMV 3' forward | GATCgaattcATTGTGTGGCAAGCTATTAC | *EcoR*I |
| | PMV 3' reverse | GATCggcgccGGGCATTTAGATTCTATAAA | *Kas*I |
| **Guide RNA/DNA and primers for confirmation of integration** | | | |
| gRNA | | ATAATGTTGAAGATATT/GTG*TGG* | |
| gDNA | | TAAGTATATAATATTATAATGTTGAAGATATTGTGGTTTTAGAGCTAGAA | |
| Confirm integration | *a* | TTCATCTTCGTTAAGTTTCCCGTGTAATGG | |
| | *b* | CAACATTATTAGATACAATTACATCTTTTGAATTATTTTCCTCATC | |
| | *c* | ATACGCATAATCGGGCACATCATAGGGATA | |
